# Nuclear translocation triggered at the onset of hearing in cochlear inner hair cells of rats

**DOI:** 10.1101/2022.03.04.483048

**Authors:** Megana R. Iyer, Radha Kalluri

## Abstract

Nuclear position is precisely orchestrated during cell division, migration, and maturation of cells and tissues. Here we report a previously unrecognized, programmed movement of the nucleus in rat cochlear inner hair cells. This cell-type specific nuclear movement among hair cells of the inner ear offers a new glimpse into the cellular mechanisms involved in the functional maturation of inner hair cells. In early post-natal days, the nucleus sits at the very basal pole of the hair cell, far from the apically located mechano-transducing stereocilia, but close to where synapses with primary afferent and efferent neurons are forming. By maturity, the nucleus moves to occupy a new position half-way along the length of the cell. Remarkably, nuclear translocation happens quickly over the course of 1-2 post-natal days, coinciding with the onset of hearing. The movement of the nucleus likely signals the closing of a critical period correlated with the forming and refining of synapses at the onset of hearing.

**Significance Statement:** The misplacement of a cell’s nucleus has been implicated in hearing, vision, and muscular pathologies. In this study, we report a remarkably abrupt change in nuclear position in cochlear inner hair cells that coincides with the onset of hearing in rats, when hair cells switch from producing sensory-independent action potentials to sensory-dependent graded potentials. This work suggests that post-mitotic nuclear migration may be critical to the final phase of post-natal development in mammalian hearing.

## Introduction

The position of a cell’s nucleus is important for supporting various cellular functions. The precision with which the nucleus is maneuvered in polarizing, migrating and mitotic cells highlights the importance of nuclear position at different points in a cell’s development. For example, in developing neuroepithelia, nuclei move between the apical and basal poles of the cell during each phase of the cell division cycle in a process called interkinetic nuclear migration (Del Bene 2011). The apical position during division ensures that the two daughter cells each receive a nucleus and properly integrate into the rest of the tissue (Gundersen and Worman 2013). Disruption of this process leads to unbalanced cell fate decisions, aberrant differentiation (Schenk et al. 2009), and loss of epithelial structure (Spear and Erickson 2012).

Emerging evidence from several systems suggests that nuclear position is also important in differentiated post-mitotic cells. Hearing, vision, and muscular pathologies have all been noted from misplaced nuclei (reviewed in Razafsky and Hodzic 2015). In the outer hair cells of the mammalian auditory system, mutations that disrupt the links between the nuclear envelope and cytoskeleton cause nuclear misplacement leading to cell degeneration and deafness (Horn et al. 2013). Similarly, in cone photoreceptors, the nucleus moves along the basal-apical axis of the cell during the final phases of post-natal development (Rich et al. 1997). Disruption of the **Li**nkers of **N**ucleoskeleton to **C**ytoskeleton (LINC) complex prevents this movement resulting in reduced synaptic efficiency and retinal dysfunction (Xue et al. 2020).

The reason why nuclear position matters for these cells remains unclear. One possibility is that the influence of extrinsic signals on nuclear transcription is regulated by nuclear position. Alternatively, genetic programs responding to developmental cues may direct construction of specialized cellular structures such as the ribbon synapse and mechano-transduction machinery. Accurate construction of these structures may require the careful positioning of the nucleus within the cell. For instance, in multi-nucleated muscle cells, nuclei are normally positioned at the peripheries of the cell and close to synapses. The proximity to synapses appears to promote appropriate synaptogenesis as nuclear mispositioning results in improper synaptic development and muscle dysfunction (Razafsky and Hodzic 2015). Although the details of how transcription relates to nuclear position remains to be understood, extrinsic signals are known to be transduced into the nucleus. For example, the mechanical forces that push and pull the nucleus are known to be transduced via the cytoskeleton through the nuclear envelope (reviewed in Graham and Burridge 2016; Jahed and Mofrad 2019) to influence the chromatin structure and organization of the genome (Uhler and Shivashankar 2017; Lammerding and Kirby 2018)

Here we describe a post-mitotic nuclear migration in the inner hair cells of the mammalian cochlea. This movement is remarkably abrupt and coincides with the onset of hearing, when inner hair cells’ physiology transforms from spiking (i.e. producing action potentials) to non-spiking (reviewed in Kros 2007). In contrast, nuclear position does not change in outer hair cells, whose nuclei remain polarized to the base of the hair cell throughout post-natal development. Using immuno-histochemistry, we characterized the timeline of nuclear migration and compare this to previously reported changes in inner hair cell morphology and physiology. We propose that nuclear migration and the next phase of hair cell development are linked and likely triggered by the larger transduction currents triggered by the onset of hearing.

## Methods

All procedures were approved by the animal care and use committee at the University of Southern California. Chemicals were obtained from Sigma-Aldrich (St. Louis, MO), unless otherwise specified.

### Animals

All data in this paper were collected from Long-Evans rats ranging in age from postnatal day (P)3-69. All animals were handled and housed in accordance with National Institutes of Health *Guide for the Care and Use of Laboratory Animals.* All animal procedures were approved by the University of Southern California Institutional Animal Care and Use Committee.

### Cochlear dissection and immunostaining

Cochlear samples were prepared from rats ranging in age from P3 through P69. Otic capsules were dissected in chilled and oxygenated Leibovitz-15 (L-15) medium supplemented with 10 mM HEPES (pH 7.4, ~315 mmol/kg). Samples were fixed in 4% PFA for 60-90 minutes at room temperature. At older ages (P18 and above), samples were decalcified in 10% EDTA for 2-5 days before fine dissection. Between 3-7 cochleae contributed to the data set at most ages.

Fixed cochleae were transferred to phosphate-buffered saline (1×PBS, pH 7.4). Fine dissection began by removing the bony covering of the otic capsule to reveal three turns of the cochlea. Sharp scissors were used to separate the apical, middle, and basal turns. The stria vascularis, excess bone, and other overlaying membranous layers including the tectorial membrane were removed using fine forceps. Turns were incubated in a blocking buffer containing 16% normal goat serum, 0.03% Triton X-100, 120 mM phosphate buffer, and 450 mM NaCl for 1 hour at room temperature. After three rounds of wash in 1x PBS for 5 minutes each, preparations were incubated in dilutions of primary antibodies at room temperature for 16-20 h or at 4°C for 72 h. Secondary antibodies were then added and incubated for one hour at room temperature. Following the primary and secondary incubations, we washed the samples for three 10 min rounds in 1× PBS.

The following primary antibodies were used to label the following structures: (1) anti-CtBP2 mouse monoclonal IgG (Becton Dickinson Company) to label the synaptic ribbon-specific protein RIBEYE, (2) anti-myosin-VI-rabbit poly clonal (Proteus Bioscience Inc.) to label the cytoplasm of inner hair cells, and in select samples, (3) anti-beta III tubulin mouse IgG2a (TUJ1; Covance) to label cochlear neurons. The following secondary antibodies acquired from Life Sciences Inc. were used: Alexa Fluor 488 anti-mouse and Alexa Fluor 594 anti-rabbit. Samples at different ages were typically processed on different days when animals of the correct age became available. Samples from several experiment days were pooled to provide the data at any particular age.

A subset of measurements was made from cochlear scans collected as a part of another project. The goal of those experiments was to record from spiral ganglion neurons (see Markowitz and Kalluri 2020 for details). As such, the samples were not immediately fixed and the cochleae were first exposed to an enzymatic cocktail comprised of trypsin and collagenase in L-15. Hair-cell morphology was not noticeably different between the age matched samples collected for the present purpose and those that were reanalyzed from the previous data set. However, wherever possible, these data are distinguished by a different symbol.

### Imaging

Fixed and labeled samples were mounted under a glass coverslip and onto glass slides with the fade-protectant medium Vecta-shield (Vector Laboratories). Hardened nail polish dots were placed under the coverslip to prevent it from crushing the cochlear sections. We generated a series of z-stack images of immunofluorescent signals using one of the following two confocal microscopes (1) Olympus FV1000 laser scanning confocal microscope and (2) Zeiss LSM 800 with a 40-63x, 1.42 numerical aperture, and an oil immersion objective. To visualize the entire inner hair cell, z-stack images were taken from the cuticular plate to the very basal pole of the cell. Each channel was scanned in sequence with the same optical thickness. Laser power, gain, and offset settings for each channel were set to optimize the dynamic range of the fluorescent signal in each sample (between 0 and 4095).

### Analysis

Analysis was done on two sets of data: the first set of images were taken from confocal scans of samples that were fixed after electrophysiological recordings (Markowitz and Kalluri 2020), while the second were fixed immediately after euthanasia.

Both length calculations and the analysis of nuclear position in hair cells were done offline using Icy (http://icy.bioimageanalysis.org/). Images included in this paper were processed using Imaris software to produce 3D projections of samples.

Measurements were taken primarily in the XY and XZ planes. The 3-D region of interest tool was used to place a point at the end of the cuticular plate, at the top end of the nucleus closest to the basal pole, and at the basal-most region of the inner hair cell (see arrows in Figure 1, A2). For hair cells that appeared bent, a fourth point was placed at the base of the nucleus, closest to the cuticular plate, in order to accurately capture the full length of the cell.

**Figure 1.**
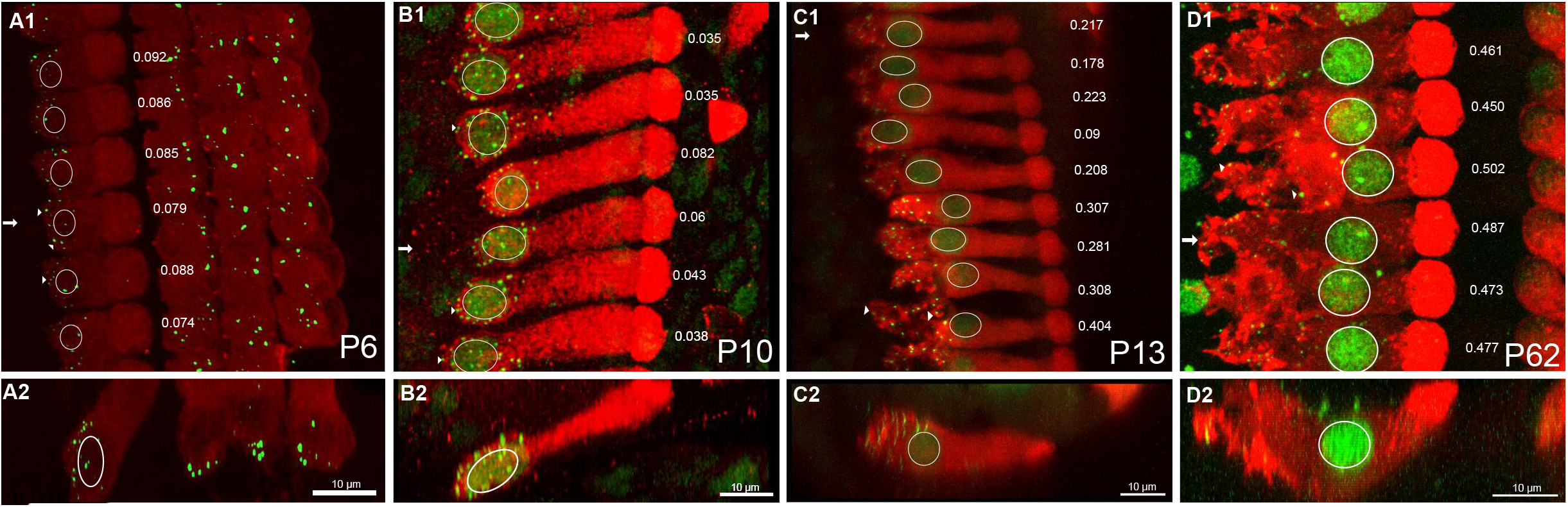
A dramatic change in nuclear position and hair cell morphology occurs around the onset of hearing. **A1-D1.** Confocal images of inner hair cells in the middle cochlear turn at P6, P10, P13, and P62 in the XY plane. Inner hair cells were labeled with Myosin VI (red) and nuclei and synaptic ribbons (arrowheads) were labeled with CTBP2 (green). **A2-D2.** Same samples as above shown as a cross-section in the XZ plane to view the length of the entire inner hair cell. **A1,B1.** Before the onset of hearing, nuclei appear to be located at the base of IHCs and synaptic ribbons are clustered around the nucleus. **C1.** By P13 in the middle turn, variance in nuclear position is apparent. Ribbons have dispersed throughout the basal pole. **D1**. By P62, the nucleus has reached its final position in the middle of the cell. A full migration seems to have occurred by P62. Outer hair cells, though present and labeled, were cropped out of these scans.

Lengths were calculated using all three dimensions of each point. L1 was defined as the supranuclear length, or the length between the cuticular pole and the top of the nucleus, and L2, the subnuclear length, was defined as the length between the top of the nucleus and the basal pole. Nuclear position was calculated by dividing L2 by the total length of the hair cell (L1 + L2); here, we normalized by cell length to account for differences in length across the cochlea and with maturation. Normalizing by length was also necessary to account for fixation or mounting artifacts that may have caused stretching of the cells.

### Statistical Analysis

Welch’s ANOVA with a post-hoc Games Howell test was used to compare the following parameters: nuclear position in pre-hearing IHCs and post-hearing IHCs; and nuclear position as a function of cochlear turn at a critical period. The level of significance for this study was p < 0.05.

## Results

We imaged inner and outer hair cells (IHCs and OHCs, respectively) from the apical, middle and basal turns of cochleae in rats ranging in age from P3 to P69. Examples of labeled inner hair cells and nuclei from the middle cochlear turn are shown in the XY and XZ dimensions at P6, P10, P13, and P62 (Figure 1A, B, C and D). Nuclei and synaptic ribbons were fluorescently labeled using an antibody against c-terminal binding protein 2 (CtBP2, green). We visualized the cytoplasm and boundaries of the hair cells by fluorescently labeling with antibodies against the motor protein, Myosin VI (red).

The positions of inner hair cells’ nuclei change dramatically during the post-natal maturation. During the first two post-natal weeks (e.g., P6 and P12, Figure 1A,B), nuclei are located at the very base of IHCs, leaving little Myosin VI labeled area under the nucleus. Synaptic ribbons are found tightly clustered near the nucleus at the basal pole of the cell (arrowheads). By contrast, in mature inner hair cells, nuclei are found near the middle of the hair cell (Figure 1D) with a substantial sub-nuclear region (Myosin VI enriched). Ribbons are found at the basal periphery of the sub-nuclear region and well below the mid-line position of the nucleus (arrowheads). At P13, nuclear position is highly variable and ranges between the basal pole and middle of hair cells (Figure 1C).

The change in the nuclear position of inner hair cells is contrasted by the relative age invariance of nuclear position in outer hair cells. Nuclei in outer hair cells remain located at the bottom of the cell throughout early post-natal development and into maturity (Figure 2).

**Figure 2.**
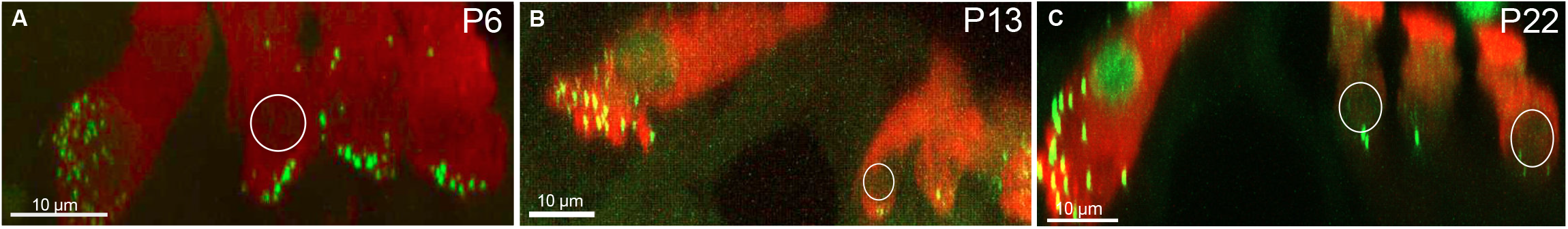
Outer hair cell nuclei remain basal bound throughout development. Outer hair cells in the middle turn are outlined at P6 **(A)**, P13 **(B)**, and P62 **(C)**. Unlike inner hair cells, nuclei in OHCs are positioned at the basal pole of the cell in all samples.

To characterize the timeline over which nuclear position changes in inner hair cells, we quantified nuclear position relative to total hair cell length as a function of post-natal age (Figure 3). To do this, we measured a series of high-resolution confocal scans through the inner hair cells (e.g., the images in Figure 1). Next, we bisected each cell using an oblique slice through its long axis (this was typically very close to a plane in the XZ dimension). Then we measured the length of the subnuclear and supranuclear regions of each cell (Figure 3A, inset). The subnuclear length, *L*_1_, was measured from the bottom of the nucleus (the point closest to the basal pole) to the basal-most point of the hair cell. Supranuclear length, *L*_2_, was measured piecewise as the sum of two lengths; the nuclear length plus the length from top of the nucleus to the cuticular plate. The sum of *L*_1_ and *L*_2_ was taken as the total cell length. Note that the piece-wise measurement was effective at capturing cell length even in samples where the hair cell appeared to bend (e.g. Fig. 1D.3). In addition to raw lengths, we computed nuclear position relative to the total length of the cell: 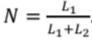. The relative position takes into account sample to sample variations in cell length that might arise due to maturation or methodological variability.

**Figure 3.**
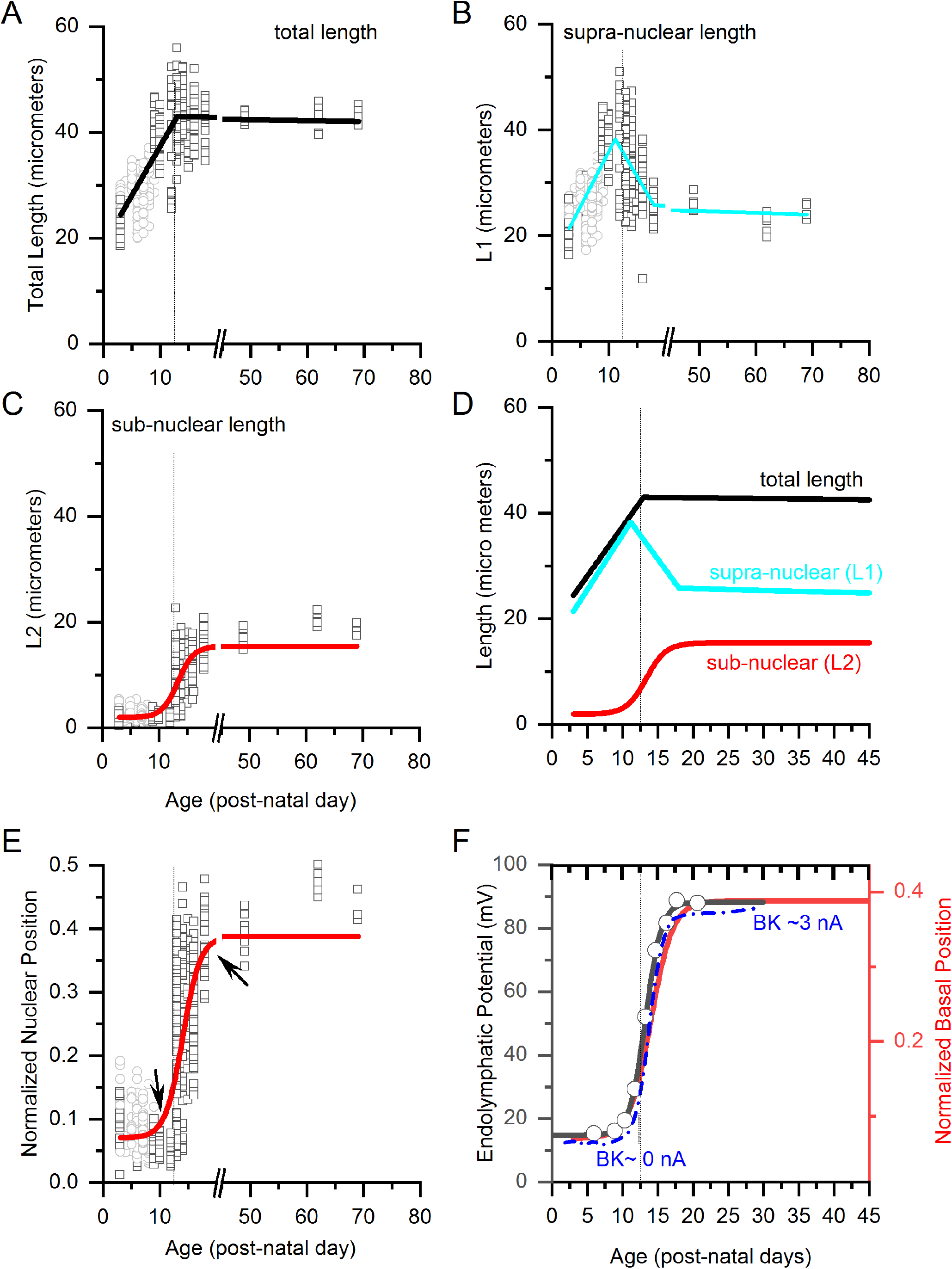
Quantification of nuclear position reveals a period of progressive cell growth followed by an abrupt nuclear translocation event at the onset of hearing. **A.** Total IHC length as a function of post-natal age ranging from P3 to P69 pooled across all turns. Open squares are length measurements from individual hair cells from samples in which cochleae were fixed immediately after euthanasia for immunohistochemistry. Open circles indicate samples in which samples were fixed after electrophysiological recordings (Markowitz and Kalluri 2020).Cell length grows progressively until it reaches a maximum around P13. **B.** Supra-nuclear length (*L*1) increases progressively as a function of age until the onset of hearing when it suddenly begins to decrease. **C**. Sub-nuclear length is relatively small until about P13 when it begins to increase. **D.** The trendlines describing total (black), supra-nuclear (cyan), and sub-nuclear lengths (red) are drawn on the same axis to compare the timelines for age dependent changes in lengths. The coincidence between the decrease in supra-nuclear length with the increase in sub-nuclear length indicates that the nucleus is moving. **E.** Relative nuclear position, which characterizes nuclear position while accounting for length changes (*L*_2_ / (*L*_1_ + *L*_2_)), increases suddenly around P13. **F.** The emergence of EP (black curve and points, reproduced from Bosher and Warren 1971) in rat IHCs coincides with the onset of nuclear translocation (red curve, y-axis to the right); both events are abrupt and occur just at the onset of hearing. The emergence of BK channel currents (blue curve, reproduced from Kros 1998). The age axis for EP data was adjusted from the originally reported gestational age to post-natal day. Note that the BK current data are from mouse, whereas EP and nuclear position data are from rat. The inset schematic in (A) shows how different lengths were measured. The dashed line in all plots is drawn at P13. Trend lines were drawn either by piecewise linear fitting (A&B) or by a Boltzmann fit (C&D).

### Nuclear position changes because nuclei move and not because hair cells grow

Total hair cell length increases progressively as a function of age during the first two post-natal weeks, at which point it reaches a maximum and length becomes relatively age-invariant (Figure 3A). A maturational increase in cell length is consistent with previous electrophysiological reports showing that hair cell capacitance increases with post-natal age (Corns et al. 2018).

As we noted in Figure 1, nuclear position changes dramatically before and after the onset of hearing. Given that cell length changes during maturation, changes in the relative position of the nucleus from the basal pole to the middle of the cell (e.g. Figure 1) could result from either an outgrowth of the sub-nuclear region and/or a migration of the nucleus. To determine which of these scenarios is at play, we examined age-dependent changes in the sub-nuclear and supra-nuclear lengths of the inner hair cells (Figure 3B and 3C). Much like total cell length, supra-nuclear length grows progressively with age, reaching a maximum around P14 (Figure 3B). During this period, sub-nuclear lengths are small and do not change (Figure 3C). In other words, the nucleus is basal-bound while the cell grows. Then, around P14, an abrupt change occurs, and sub-nuclear lengths suddenly begin to increase, while supra-nuclear lengths decrease. Note that by this time the cell length has stopped changing (compare the time course for total, sub-nuclear, and supra-nuclear changes in Figure 3D). The simultaneous increase in sub-nuclear length coupled with a decrease in supra-nuclear lengths, while total length stays constant, indicates that the nucleus is moving from the bottom to the middle of the cell.

### Change in nuclear position in inner hair cells coincides with the onset of hearing

The onset of nuclear migration is remarkably abrupt, beginning around P13 and completing by P16. Note that the onset of migration is much more abrupt than that captured by the Boltzmann fit in Figure 3E. Remarkably, the moment at which nuclear position abruptly changes coincides with the rapid emergence of endolymphatic potential (EP; Figure 3F, data from Bosher and Warren 1970) and the subsequent onset of acoustic hearing in rats. The coincidence in the timing between onset of EP and nuclear translocation suggests that the larger transduction currents triggered by the onset of hearing contribute to the movement of the nucleus. That nuclei are migrating is especially evident when comparing adjacent cells in individual cochlear turns within the P13 through P14 age range (e.g., Fig. 1C.2). Here, note that adjacent hair cells of similar lengths in the middle turn have nuclei at very different positions (also see Figure 4). This variance in nuclear position indicates that the image caught cells at different stages of a nuclear translocation.

**Figure 4.**
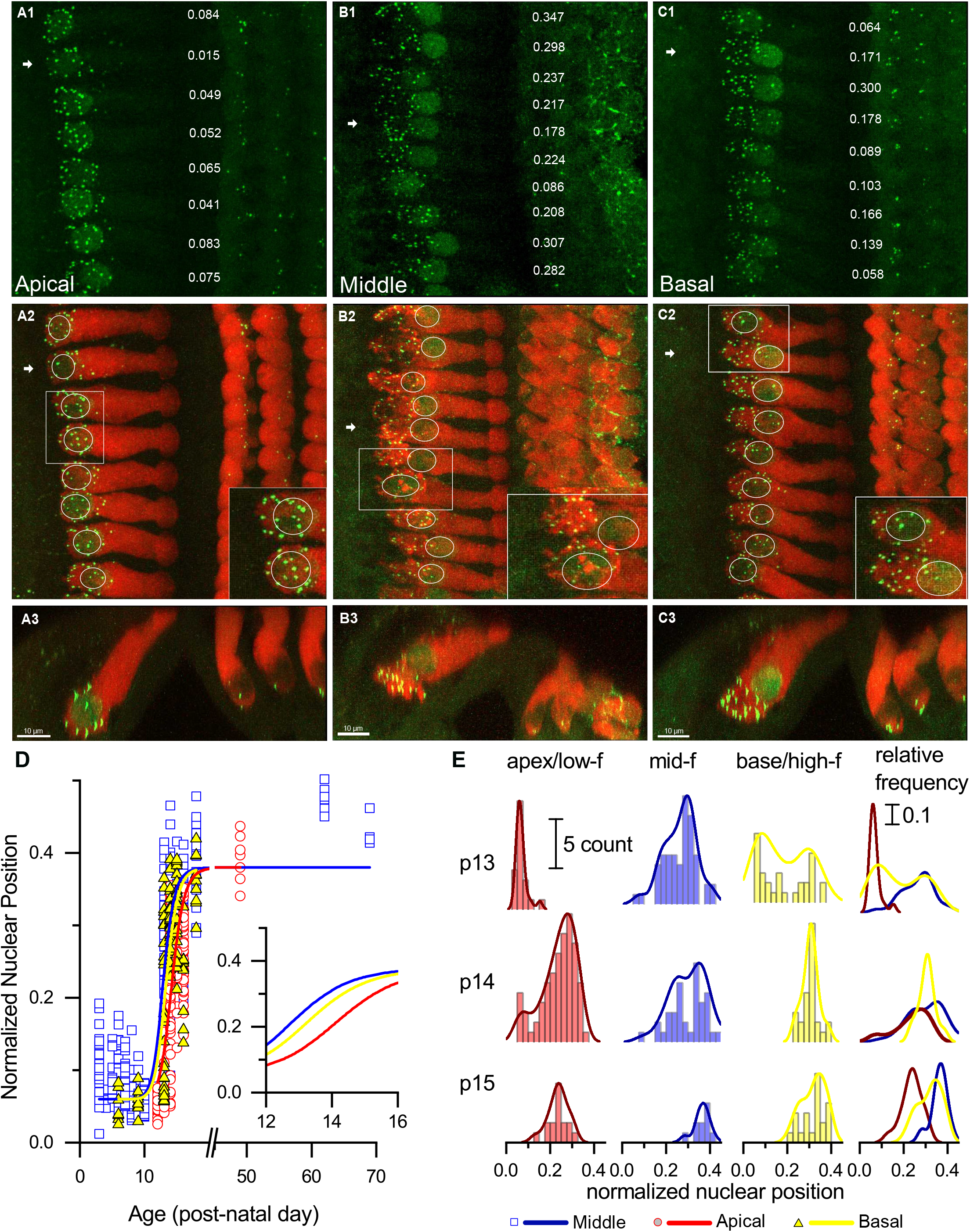
Nuclear translocation happens first in the middle and basal turns and then in the apical turn. **A1-C1.** CTBP2 labeled nuclei and ribbons in apical, middle, and basal turns from the same P13 sample. **A2-C2.** Merged images of inner hair cells labeled with Myosin VI (red) and CTBP2 (green) in the XY plane. In the apical turn, ribbons are clustered around the nucleus, which is bound to the base of the hair cell. Nuclear position is variable in the middle turn, and ribbons are dispersed across the basal pole in both the middle and basal turns. **A3-C3.** Images of all three turns in the XZ plane. Here, outer hair cells are visible and nuclei were located at the base of cells in all three turns. At P13, most nuclei in the apical turn inner hair cells are found at the base of the cell. In contrast, nuclear position is more variable in the middle and basal turns. Note the instances of adjacent cells with similar lengths but different nuclear positions (insets). Variance in nuclear position across adjacent cells suggests that this sample has hair cells caught in the middle of transitioning between two distinct developmental states. **D.** Relative nuclear position as a function of age for all three cochlear turns from P3-P69. Nuclear migration in the middle turn begins earliest, followed by migration in the basal and apical turns, respectively. Nuclear migration in the apical turn lags 1-2 days behind the middle turn. **E.** Histograms showing relative nuclear position in apical (red), middle (blue), and basal (yellow) turns at P13, P14, and P15. Smoothed distributions are overlaid on top of each histogram and compared on the same axis. Note that the distributions are largely bimodal with one mode at approximately 0.07 and another at 0.4. The heights of the two modes indicate the relative developmental state of the groups of hair cells; with apical hair cells lagging the basal turn and middle turns.

In summary, these results indicate that after a period of progressive cell growth, the inner hair cell suddenly stops growing and the nucleus migrates from the basal pole to the middle of the hair cell.

### Nuclear migration occurs first in the middle cochlear turn

Next, we examined the time course of nuclear translocation across the cochlear length. To do this we sorted the data shown in Figure 3 according to one of three cochlear turns; the low-frequency encoding cochlear apical turn, the mid-frequency encoding cochlear middle turn, and high-frequency encoding cochlear basal turn. Before P12, nuclei appeared to be positioned at the bottom/basal pole of the inner hair cell in all three turns. By P13, IHCs in the middle turn showed signs of a migration event (compare the nuclear positions in Figure 4 A,B,C, see also the insets). Nuclei in the apical turn, however, are almost exclusively found at the basal pole of the inner hair cell (Figure 4A). Note the striking variability in nuclear positions in the three cochlear turns of the same cochlea (Figure 4 B, C).

When compared to the observed timeline in the middle turn, the onset of nuclear translocation is delayed in apical and basal turn cochleae, as illustrated in Figures 4 D & E. In 4D, the relative nuclear position across age is shown with the data now color coded to indicate cochlear turn. The Boltzmann curves through the nuclear position data are shifted to the right for the basal and apical turns when compared to the middle turn.

To further quantify differences across cochlear turn, Figure 4E shows the distribution histograms of nuclear position at P13 and P14. The smoothed distribution curves that are overlaid on the histograms are largely bimodal, with one mode at the bottom of the hair cell (nuclear positions around 0.07) and the second mode near the middle of hair cells (nuclear positions around 0.4). The relative height of each mode indicates the proportion of nuclei that are in either the immature position at the bottom of the hair cell or the mature position in the middle of the hair cell. In the apical turn at P13, most of the inner hair cells nuclei are in first mode at the immature position. In the middle turn, many nuclei are already in the mature second mode. The basal turn is more advanced than the apical turn in having some nuclei that have moved to the mature position but perhaps not as advanced as the middle turn because many nuclei are still in the immature position.

By P14 and P15 nuclear position of both the middle and apical turn hair cells has progressed, with most nuclei in the apical turn now occupying the second mode, although nuclear position is still significantly different between middle and apical turns ( F(1, 58) = 26, p < 0.0001, Welch’s t-test).

## Discussion

### Nuclear translocation demarcates the transition between two distinct periods of post-natal development in inner hair cells

Here we showed that the nuclei of inner hair cells experience a remarkable sudden and timed migration from the bottom of the cell to the middle of the cell. This movement is notable in its coincidence with the onset of hearing when these cells also experience a dramatic biophysical and functional transformation. Prior to the onset of hearing, inner hair cells produce sensory-independent regenerative action potentials (e.g., Kros 1998). The early activity impacts cell growth, synaptogenesis, and the formation of upstream neuronal circuits (Jones et al. 2007; Kandler et al. 2009; Tritsch and Bergles 2010). Around the onset of hearing, inner hair cells cease to produce regenerative spikes and start responding with graded potentials that are useful for encoding the amplitude and timing of incoming sounds with high resolution. The correlation between the timing of nuclear migration and the functional transformation of the hair cells suggests that the two phenomena are linked, although the nature of the link remains to be understood.

The transformation from spiking to non-spiking behavior in inner hair cells is driven by changes in ion channel properties, including the loss of calcium channels and the acquisition of new potassium channels, such as big-conductance calcium dependent potassium (BK) channels (Kros 1998; Marcotti et al. 2003). The time course for BK channel acquisition is similar to that of nuclear translocation and happens rapidly at the onset of hearing. In contrast, other hair cell features, like cell size, increase slowly and progressively during the pre-hearing period (Kros 1998). The coincidence between changes in ion channel expression and nuclear migration suggests that the large transduction currents produced at the onset of hearing are at least indirectly triggering both changes, although the nature of the connection between nuclear migration and ion channel expression remains to be determined.

The idea that nuclear migration and the biophysical maturation of inner hair cells are linked is reinforced in animals with thyroid hormone mutations. In mice, thyroid hormone (TH) is expressed in IHCs between embryonic day (E)18 and P12. The inner hair cells in TH knockout animals develop normally until about P12 but remain immature well past the onset of hearing. The immature phenotype includes a failure or delay in acquiring BK channels (Rüsch et al. 2001), persistence of efferent connections to inner hair cells and immature looking synapses (Sendin et al. 2007; Sundaresan et al. 2016). It is now also apparent that the nucleus itself remains positioned to the base of the cell at P14 whereas in the wild type mice, the nucleus migrates away from the basal pole to the middle of the cell (Sendin et al. 2007 Figure 2; Sundaresan et al., 2016, Figure 7). The data from the TH knockout mice suggest that nuclear migration and the biophysical maturation of inner hair cells are connected since disruptions to one coexist with dysfunction in the other.

We hypothesize that nuclear repositioning is triggered by the onset of endolymphatic potential (EP) between P13 and P14 in rat (Bosher and Warren 1971). EP emerges first in the middle turn of the cochlea where we also first see nuclear position shifting. This idea is also consistent with the joint disruption to nucleus translocation and failure/delay in BK channel acquisition in hypothyroidism which disrupts mechano-transduction, in part, by its influence on endolymphatic potential (Rüsch et al. 2001). Similarly, hair cells fail to mature and return to an immature state in mature hair cells from mouse models with targeted disruptions to mechano-transduction (Nist-Lund et al. 2019, Geller et al. 2009, Corns et al. 2018).

#### Developmental gradients across the length of the cochlea

Nuclear migration occurred in all three cochlear turns, but this did not occur in synchrony. Nuclei began to move in the middle turn first, followed by the high-frequency-encoding basal turn and low-frequency-encoding apical turn. This wave is similar to the developmental waves seen at the very onset of hair cell development. In the mouse cochlea, progenitors in the basal turn differentiate into hair cells and supporting at embryonic day (E)13.5 (Chen et al. 2002). This wave of differentiation continues towards the cochlear apex over the following few days (Chen et al. 2002; Groves and Fekete 2012). Similarly, transduction currents mature first in the mid-basal region at P0 to reach maturity in the apex several days later (Lelli et al. 2009). If a change in nuclear position marks a new stage in hair cell development, then, the maturational changes triggered at the onset of hearing also follow a base to apex spatial gradient.

Nuclear position did not change in the outer hair cells. The stability of nuclear position in outer hair cells stands in stark contrast to the mobility of inner hair cell nuclei. Indeed, as we show here, nuclear migration is part of the natural course of post-natal development for inner hair cell whereas keeping the nucleus at the base of the cell appears to be crucial for the normal function of outer hair cells (Horn et al. 2013; Taiber et al. 2020). In mouse models with mutations to the proteins that connect the nucleo-plasma to the cyto-skeleton (**Li**nkers of **N**ucleoskeleton to **C**ytoskeleton, LINC complex), OHC nuclei float to the apex of outer hair-cells. These cells eventually die, resulting in deafness, although the reason for why the normal extreme basal position of nuclei in OHCs is significant is unclear. Curiously, the same LINC complex mutations that disrupt OHC nuclei appear to have only a modest influence on the position of inner hair cell nuclei (Horn et al. 2013), suggesting that the mechanisms controlling nuclear position may be different, even in two closely related cell types.

### Why might nuclear position matter?

Nuclear proximity to synapses may be favorable for synaptogenesis before the onset of hearing, perhaps by making it easier to deliver the components needed for forming synapses and for efficient recursive signaling between synapses and nucleus. Emerging research in skeletal muscle suggest that this might be the case for the successful formation of the neuromuscular junction (reviewed in Rasafsky and Hodzic 2015). There, the nuclei of multi-nucleated myocytes are conspicuously positioned close to the developing neuro-muscular junctions (Grady RM et al. 2005). Mutations that disrupt this proximity often lead to synaptic malformations leading to muscle pathologies (e.g., Mejat et al. 2008; Chapman et al. 2014). In the inner ear, the proximity of the nucleus to synapses in early post-natal days (when both efferent and afferent synapses are refined; reviewed in Simmons 2002) may similarly support a period of heightened synaptic plasticity.

The movement of the nucleus away from the synapses may not only indicate the end of a period of plasticity for the synapses, rather, mature function may rely on the repositioning of the nucleus. In cone photoreceptors, nuclei are first positioned near forming synapses and then migrate away from the synaptic region towards the top of cell where the photosensitive outer segment are found (Rich et al. 1997). In these cells, the efficiency of mature functioning synapses depends on the nucleus moving away from the synapse. When LINC complexes are disrupted and nuclear re-positioning does not occur (Rasafsky and Hodzic 2015), cone synapses have inefficient synaptic transmission and reduced dark sensitivity (Xue et al. 2020). Perhaps the re-positioning of the nucleus and the subsequent development of an extensive membranous and mitochondrial network between the nucleus and ribbon synapses is important for synaptic efficiency (Bullen et al. 2015). This suggests that while nucleo-synaptic proximity might be important for synaptogenesis, synaptic efficiency may require the nucleus to move away.

That the nucleus serves as an important marker for cellular sub-compartments has long been implicitly recognized in inner hair cells. Functionally distinct regions of inner hair cells are described by reference to the nucleus. For example, the ‘sub-nuclear’ region is distinct from the supra-nuclear region in cytoplasmic composition. Synapses with dense-core ribbons, endoplasmic reticulum, intracellular networks of membranes, and mitochondria are prominently found in the ‘sub-nuclear’ region (Bullen et al. 2015). On the other hand, BK channels are confined to the supra-nuclear or ‘neck’ region of inner hair cells (Pyott et al. 2004). This compartmentalization away from the synaptic region may be important for separating the synaptic and extra-synaptic regions to support the different roles of calcium in the mature cell (Pyott et al. 2004). Given that BK channels are placed with such precision to the supra-nuclear region, the position of the nucleus maybe an important cue for constraining the expression of ion channels to specific sub-cellular compartments.

Precise control of nuclear position is increasingly being recognized as important for the normal development and function of a wide range of cells (Rasafsky and Hodzic 2015). Why nuclear position matters for these cells is not well understood but a prevalent hypothesis is that nuclear position plays a role in synaptogenesis and synaptic efficiency (Xue et al. 2020). Here we described a previously unrecognized late-phase and rapid change in nuclear position in mammalian inner hair cells. The coincidence of the migration with the onset of hearing implicates nuclear repositioning as being an important step for the final maturation of these cells. Whether this rapid onset of nuclear movement is responsive to or causally related to the functional transformation of inner hair cells and their synapses remains to be determined.

## Acknowledgements

We acknowledge Maya Monges Aviles and Risha Annamaraju for technical assistance with the immunohistochemistry, imaging, and data analysis protocols. We thank Drs. Ruth Anne Eatock, Neil Segil, Daniel Bronson and Alex Markowitz for their comments on earlier versions of this manuscript.

## Author Contributions

Megana Rajam Iyer, Data curation, Formal analysis, Validation, Investigation, Visualization, Writing - original draft; Radha Kalluri, Conceptualization, Resources, Data curation, Software, Formal analysis, Supervision, Funding acquisition, Validation, Investigation, Visualization, Methodology, Project administration, Writing - review and editing.

